# Vinculin organises apical Transcellular Actin Fibres to coordinate collective migration

**DOI:** 10.64898/2026.05.29.728702

**Authors:** John James, Alexis M. Gautreau, Stéphane Romero

**Affiliations:** Laboratory of Structural Biology of the Cell (BIOC), CNRS UMR7654, École Polytechnique, Institut Polytechnique de Paris, 91120 Palaiseau, France

**Keywords:** Collective cell migration, Adherens Junctions, Vinculin, Transcellular Actin Fibres (TAFs), Arp2/3 complex, Cell unjamming

## Abstract

Collective cell migration is coordinated by adherens junctions (AJs) which serve both as structural links between the actin cytoskeleton of adjacent cells, as well as mechano-transductory structures allowing cells to transmit mechanical signals. Vinculin contributes to AJ maturation in multiple ways, binding to actin and the cadherin-catenin complex, bundling actin filaments, antagonising branched actin polymerisation and recruiting late AJ proteins. Here we have analysed the effect of vinculin on junctional actin organisation and its role in collective migration during unjamming in the human epithelial cell line MCF10A. At the apical surface of MCF10A monolayers, we found transcellular actin fibres (TAFs) that are directionally coordinated across long ranges, up to 10 cells. These TAFs are contractile and “cross” cell-cell contacts at AJs. Analysis of a vinculin knockout cell line revealed that this protein is essential for the coordination of TAFs across multiple cells. Arp2/3 activity must be tightly regulated to establish a long-range network of TAFs, since its downregulation by CK666 treatment, as well as its upregulation by the expression of an activated Rac1 mutant or a mutation that prevents the vinculin-Arp2/3 interaction, all impair TAF formation. During hypotonic unjamming of monolayers, we found that MCF10A cells with long-range TAFs migrate more collectively than vinculin KO cells in which TAFs only connect adjacent cells. Similarly, space-induced unjamming of MCF10A and vinculin knockout monolayers showed that cells connected by the long-range TAFs can collectively coordinate the direction in which they extend their lamellipodia. Thus, we show that vinculin plays a novel role in organising long-range actin networks across multiple cells and coordinating collective migration within cell monolayers.

## Introduction

Collective migration is an important process in development as well as tissue maintenance. The process by which relatively static cells in mature tissues initiate migration is called unjamming, and it occurs in physiological process ranging from the healing of a wound to metastasis of cancer cells (Park et al. 2016). While collective migration in these different contexts can be governed by various signalling factors, unjammed cells are able to coordinate their migration because they are held together by adherens junctions (AJs).

AJs are structural links between the actin cytoskeleton of adjacent cells. In epithelial cells, the extracellular domains of E-cadherin from either cell bind to each other while their intracellular domains are connected to the actin cytoskeleton through α- and β-catenin. The α-catenin head binds to E-cadherin and while its tail domain binds to actin. Together this structural unit of an AJ is called a cadherin-catenin complex. Actin architecture at the junction is dynamic, with early junctions being associated with branched actin and mature junctions with linear actin (Cavey and Lecuit 2009).

Vinculin is a mechanotransductory protein recruited to cryptic vinculin-binding-sites on α-catenin which are revealed upon acto-myosin contractility (Yao et al. 2014; Yonemura et al. 2010). Vinculin is well studied in the context cell-substrate adhesions where it binds in a similar manner to talin (Ciobanasu et al. 2014; del Rio et al. 2009) but its role at AJs is less well understood. Vinculin in the cytoplasm exists in an auto-inhibited conformation with its head domain bound to its tail and is activated when recruited to cell adhesions (Johnson and Craig 1995; Chen et al. 2005). The vinculin head domain binds talin or α-catenin and the tail domain can bind actin filaments(Menkel et al. 1994). In addition to reinforcing this link between actin and the cadherin-catenin complex, vinculin serves as a signalling hub binding to a multitude of other proteins (Carisey and Ballestrem 2011) involved in adhesion maturation as well as control of cell behaviour.

Interestingly, vinculin can modulate actin architecture in multiple ways. The vinculin tail upon binding to actin can dimerise thus bundling actin filaments (Johnson and Craig 2000; Boujemaa-Paterski et al. 2020). The vinculin tail also functions as both a capping protein (Le Clainche et al. 2010) and an actin nucleator in-vitro (Wen et al. 2009), and in cells vinculin can bind and recruit other proteins which interact with actin. Most prominent among these are Arp2/3 (DeMali et al. 2002; Chorev et al. 2014) and MENA/Vasp (Brindle et al. 1996), branched and linear actin polymerisers respectively. In particular, MENA localises to tension-bearing focal adherens junctions (FAJs), where linear fibres anchor (Oldenburg et al., 2015). Arp2/3 is recruited to AJs by vinculin and the vinculin-Arp2/3 interaction has been shown to inhibit branched actin assembly, decreasing the stability of early adherens junctions (James et al. 2025).

This multi-faceted functioning of vinculin has makes it difficult to predict how it can affect the actin cytoskeleton at mature AJs. In this paper, we will utilise a vinculin knockout (VCL-/-) in the non-transformed breast epithelial cell line MCF10A to identify the effect of vinculin on the actin cytoskeleton at AJs. Additionally, vinculin being present at both cell-substrate and cell-cell adhesions has made it difficult to isolate the effect of its role at AJs. Here, we will use unjamming assays to analyse collectivity of cell groups unbiased by the speed/efficiency of migration thus isolating the effect of vinculin at AJs on collective migration.

## Results

### Vinculin coordinates Transcellular Actin Fibres across multiple cells

To mimic the mature AJs found in tissues we allowed MCF10A monolayers to grow to a jammed state and then visualised their actin cytoskeleton. Confocal microscopy allowed us to visualise cell boundaries with actin staining from the apical to the basal poles of the monolayer (Fig 1a). At the apical pole, we found a hitherto unreported organisation of actin. MCF10A cells have bundles of actin that attach one edge of the cell to the other (Fig1b,c). These bundles are coordinated across cell boundaries and remain directionally persistent across multiple cells. We called these fibres transcellular actin fibres (TAFs).

**Figure 1:**
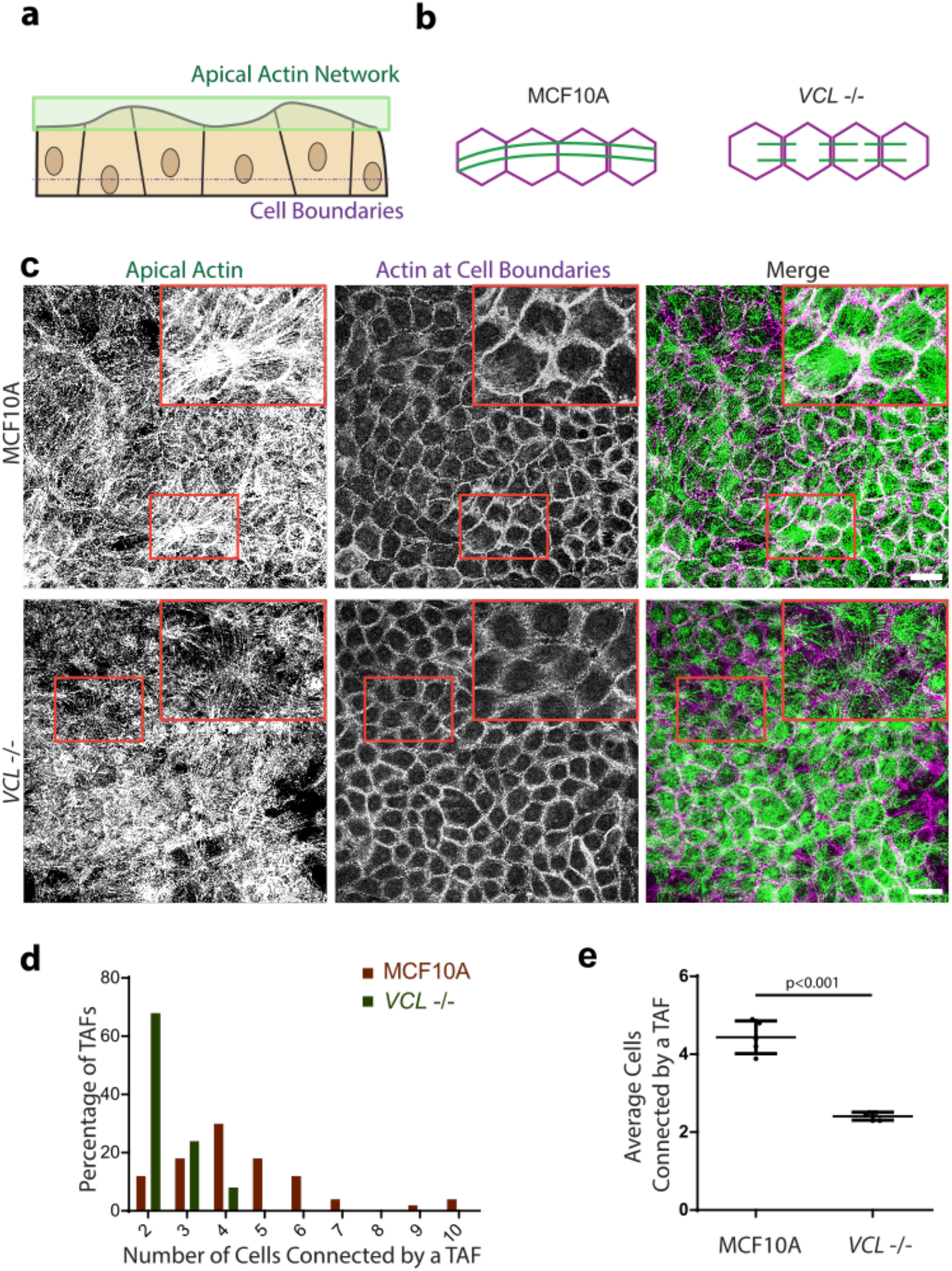
Vinculin coordinates TAFs across multiple cells. **a-c** Schematic and actin staining showing apical actin and cell boundaries in MCF10A and VCL-/-monolayers. Scale bar=20μm **d**,**e** Quantification of coordination of transcellular network. Histogram of number of cells connected by transcellular fibres and average number of cells connected by TAFs in MCF10A and VCL-/-monolayers (Mean ± SD, t-test n=5, N=3, 1 representative experiment shown.)

In VCL-/-cells, the TAFs do not directly connect one edge of the cell to another (Fig1b,c). Instead, these fibres seem to connect the cell boundary to neighbors isotropically in all directions through a disorganised actin network in the interior of cell. As a result, in VCL-/-monolayers, the TAF network seems less coordinated over long ranges. To quantify this change, we counted the number of cells a TAF connects before experiencing a change in direction of more than 30°. This is represented as a histogram of number of TAFs counted that cross a certain number of cells (Fig 1d). In MCF10A monolayers, a TAF can connect upto 10 consecutive cells with a median of 4 cells being connected. In contrast, in VCL-/-, most TAFs only connect 2 adjacent cells. On average, we found that TAFs in MCF10A monolayers connect 1.8-fold more cells compared to VCL-/-(Fig 1e). Thus, in contrast to MCF10A cells, VCL-/-form a short-range TAF network and vinculin is required to coordinate the TAF network over longer ranges.

To characterise this network further, we looked at the kinetics of formation and composition of TAFs. TAFs are not present in MCF10A monolayers 1 day after seeding cells but are well formed in mature monolayers, 3 days post seeding (Fig 2a). Unsurprisingly, vinculin is localised to sites where these fibres connect adjacent cells indicating that these fibres connect AJs at either end of the cell (Fig 2b). These AJs are distributed as puncta along the cell-cell contact and extend outwards radially from the cell. The associated actin is perpendicular to the surface of cell-cell contact indicating they are focal adherens junctions (Oldenburg et al. 2015). Further, we found that sites where several bundles converge are enriched in phospho-myosin (Fig 2c), showing that these are contractile stress fibers. Perhaps most interestingly, we found that these fibres are formed in mammary organoids developed from self-organisation of Parental MCF10A cells (Fig 2d). These organoids, called acini (Debnath et al. 2003), are a model system for mammary glands, secreting a basal membrane labelled with laminin, retaining a spherical shape and forming a lumen by cell apoptosis in the centre. Imaging the network at high resolution proved challenging in these large structures, but staining of E-cadherin revealed AJs at the surface of the acini and faint actin bundles that are coordinated across multiple cells (red arrows).

**Figure 2:**
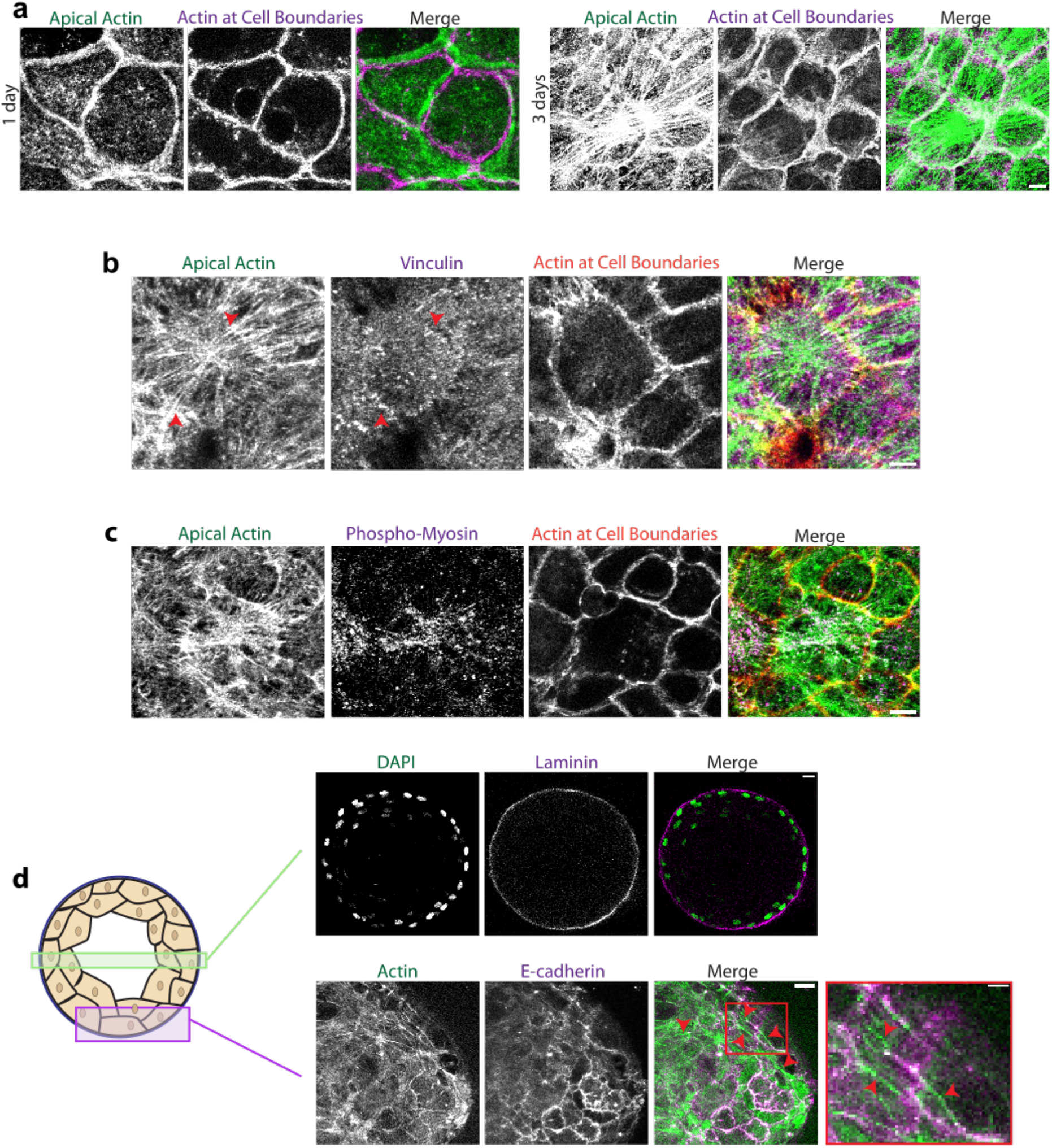
Characterization of TAFs. **a** Actin staining showing apical actin and cell boundaries in parental MCF10A monolayers 1 and 3 days post seeding. Scale bar = 5 μm **b** Staining of actin and vinculin in parental MCF10A cells. Scale bar = 5 μm **c** Staining of actin and phospho-myosin light chain 2 (Thr18/Ser19) in parental MCF10A cells. Scale bar = 5 μm **d** Staining of nuclei, basal membrane, actin and e-cadherin in MCF10A acini Scale bar = 15 μm.

### Arp2/3 activity is finely tuned to form TAFs

Considering there was a change in actin architecture at AJs, we hypothesised that the Arp2/3 complex, which is responsible for branched actin polymerisation and can associate with vinculin, could have a role in TAF formation. We grew Parental MCF10A monolayers in different concentrations of CK-666, an inhibitor of Arp2/3. We found that at low concentrations(6.25µM) of CK-666, the long-range TAF network was disrupted (Fig 3a). TAFs mostly connected 2 adjacent cells (Fig 3b) and the average number of cells connected by fibres dropped 1.7-fold (Fig 3c) compared to cells grown without CK-666. At high concentrations of CK-666 (50µM), the apical actin cytoskeleton was completely disrupted and there were no TAFs present (Fig 3a). Thus, Arp2/3 activity is required both to form TAFs and to coordinate them over long ranges.

**Figure 3:**
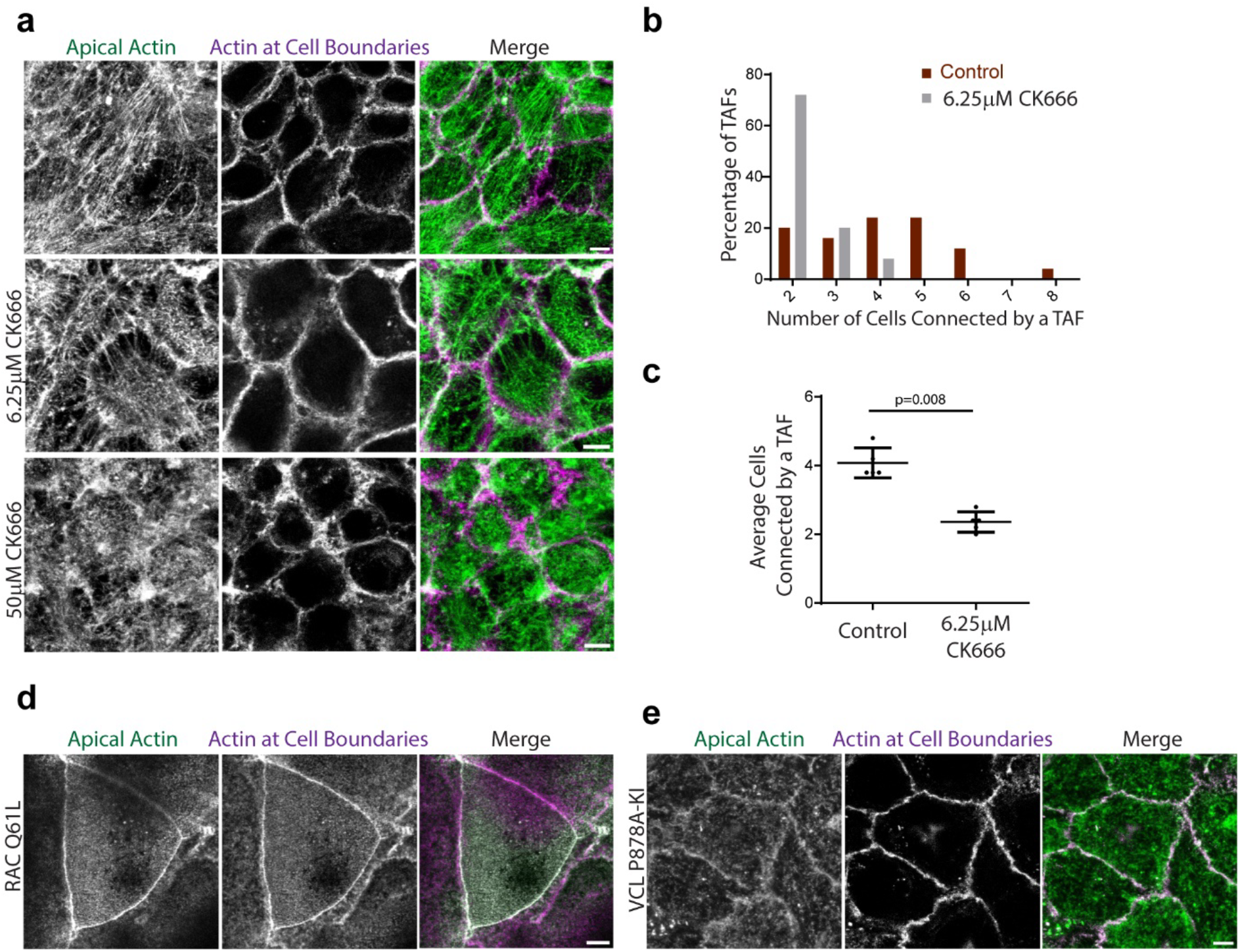
Arp2/3 activity is finely tuned to form apical actin fibres. **a** Actin staining showing apical actin and cell boundaries in MCF10A monolayers grown with and without CK666. Scale bar = 5 μm **b**,**c** Histogram of number of cells connected by transcellular fibres and average number of cells connected by TAFs. (Mean ± SD, t-test n=5, N=2, 1 representative experiment shown.) **d**,**e** Actin staining showing apical actin and cell boundaries in RAC Q61L and VCL P878A-KI cells. Scale bar = 5 μm

This finding caused us to question if increasing Arp2/3 activity would lead to formation of bundles coordinated over even longer ranges. We expressed a constitutively active mutant of the Arp2/3 activator RAC1 harboring the Q61L mutation, to constitutively activate Arp2/3 in MCF10A cells. Monolayers formed by RAC1 Q61L cells did not show any apical actin staining in the interior of the cell; only cell boundaries were visible even at the apical surface (Fig 3d). This suggests that upregulation of Arp2/3 activity also leads to impairment in formation of TAFs. Taken together, these results show that Arp2/3 activity must be finely regulated to form the transcellular actin network since both upregulation and downregulation of Arp2/3 lead to disruption of the network. Additionally, since the vinculin-Arp2/3 interaction has been shown to antagonise Arp2/3 activity in epithelial cells (James et al. 2025), one could predict that mutating the Arp2/3 binding site on vinculin would lead to a similar actin architecture compared to the RAC Q61L expressing cells. Indeed, we see in VCL P878A-KI monolayers that no TAFs are formed (Fig 3e) confirming that the regulation of Arp2/3 by its direct interaction with vinculin plays an important role in the formation of the TAF network.

### Vinculin coordinates collective migration during unjamming

To understand how this TAF network effects cell behaviour, we first looked at how it affects the environment of individual cells in the monolayer. We observed cells stained for actin and the FAJ marker MENA (Oldenburg et al. 2015) and found that in contrast to parental MCF10A cells (Fig 4a), cells in VCL-/-monolayers are connected to almost every adjacent cell by short TAFs (Fig 4c). In parental MCF10a monolayers, contractile TAFs connect cells in specific directions potentially imparting an anisotropy to the forces experienced by cells. To quantify this anisotropy, we analysed the orientation of FAJs along contractile TAFs which allow cells to exert forces on their neighbours. Fig 4b shows the orientation of all FAJs in a cell normalised to the average orientation which represents the direction of the net pulling force experienced by a cell. We found that MCF10A cells have their FAJs and associated TAFs oriented along a specific main axis, reflecting a highly anisotropic force distribution, while VCL-/-cells have FAJs oriented all around the cell, indicating an isotropic distribution of junctional tension.

**Figure 4:**
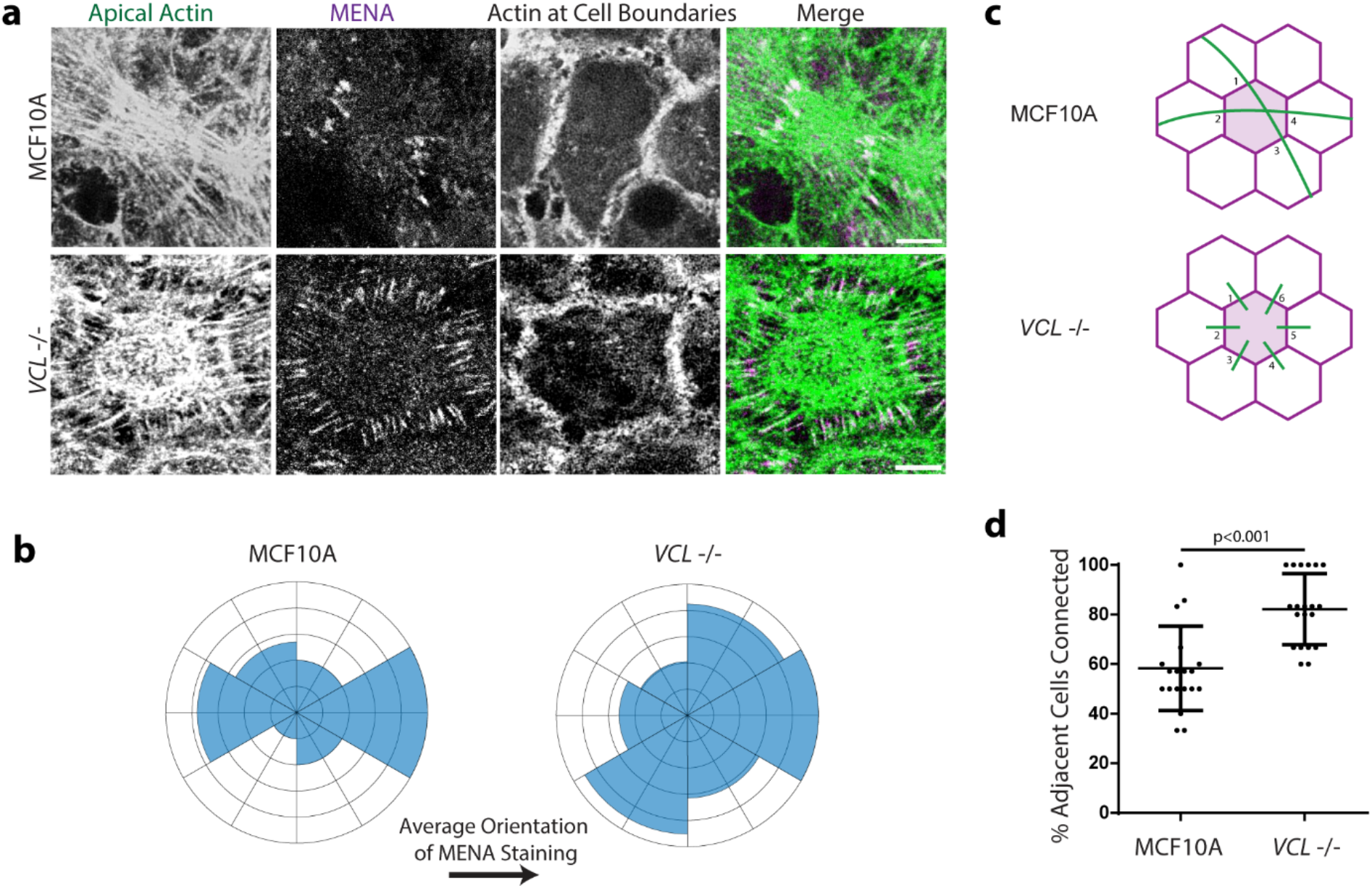
Organised TAFs create anisotropy in the monolayer. **a** Staining of actin and MENA in Parental MCF10A and VCL-/-cells. Scale bar = 5 μm **b** Orientation of FAJs labelled with MENA normalized to average orientation for each cell (n=5, N=3, 1 representative experiment shown.) **c**,**d** Schematic and quantification showing percentage of adjacent cells connected by a TAFs in MCF10A and VCL-/-monolayers (Mean ± SD, t-test n=18, N=3, 1 representative experiment shown.)

Since in-silico studies have predicted that force anisotropy in cell monolayers can lead to collective migration, we hypothesised that the TAF network would influence the collectivity of migration. We utilised hypotonic unjamming as a model for studying collective migration in monolayers (Hakim and Silberzan 2017). Cells grown in culture tend to divide and migrate to fill the space available to them and eventually stop moving. Treating such jammed cells with hypotonic conditions induces collective migration of cells confined within domains. We found that the TAF network was retained upon hypotonic unjamming (Fig 5a) and as with jammed monolayers, parental MCF10A cells had multiple cells (3.5 on average) connected by TAFs while they predominantly connected only adjacent cells in VCL-/-(Fig 5 b,c). We utilised particle image velocimetry (PIV) to plot vectors showing the direction of movement of particles in the field and found that the vectors seemed to be correlated over larger distances in MCF10A compared to VCL-/-(Fig 5d). This suggests that MCF10A cells form larger velocity domains compared to VCL-/-, and to quantify this we calculated the distance over which the vectors are correlated (Garcia et al. 2015). MCF10A cells have a larger correlation distance, around 4 cell diameters, while VCL-/-have a lower correlation distance of around 2 cell diameters (Fig 5e). Together this shows that MCF10A cells with a long-range network migrate more collectively than VCL-/-cells with a short-range network.

**Figure 5:**
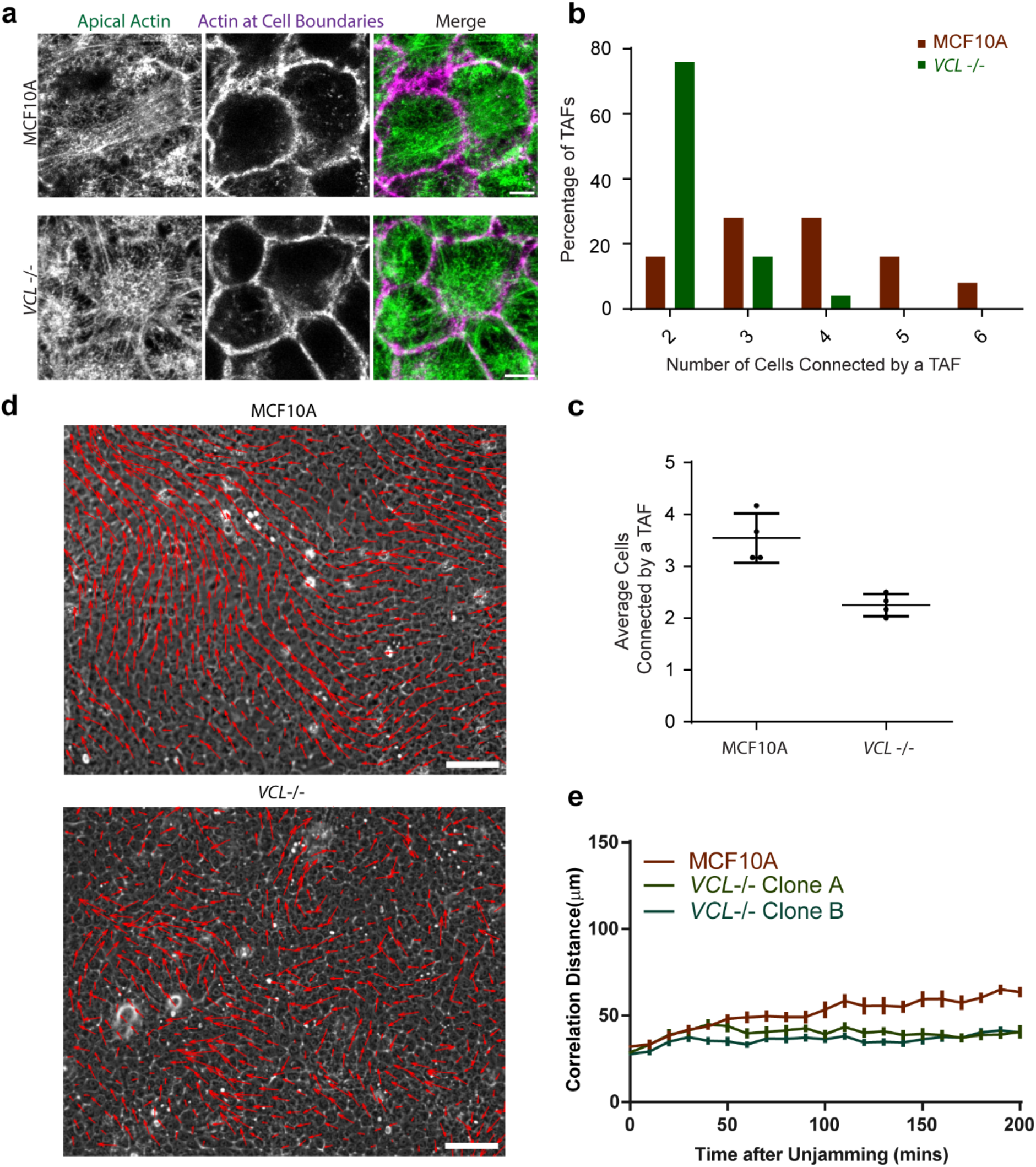
Vinculin coordinates collective migration during hypotonic unjamming. **a** Actin staining showing apical actin and cell boundaries in MCF10A monolayers 10 min after unjamming. **b**,**c** Histogram of the number of cells connected by transcellular fibres and average number of cells connected by TAFs. (Mean ± SD, t-test n=4, N=3, 1 representative experiment shown.). **d** Displacement vectors from PIV analysis overlaid on phase contrast images of MCF10A and VCL-/-monolayers after hypotonic unjamming. Scale bar = 100 μm. **e** Correlation distance of migration for Parental MCF10A and VCL-/-cells after hypotonic unjamming. (Mean ± SD n=5, N=3, 1 representative experiment shown.)

Unjamming of cells can be achieved through a mechanical stimulus as well, by placing an inert obstacle in their path, allowing cells to jam against the surface, and then removing the obstacle. Upon space-induced unjamming, we found that the TAFs connected multiple MCF10A cells at the migrating front (Fig 6a), while in VCL-/-they only connected adjacent cells. As in monolayers, these TAFs are contractile fibres decorated with phospho-myosin and ‘cross’ cell-cell contacts at FAJs marked by MENA (Fig 6b).

**Figure 6:**
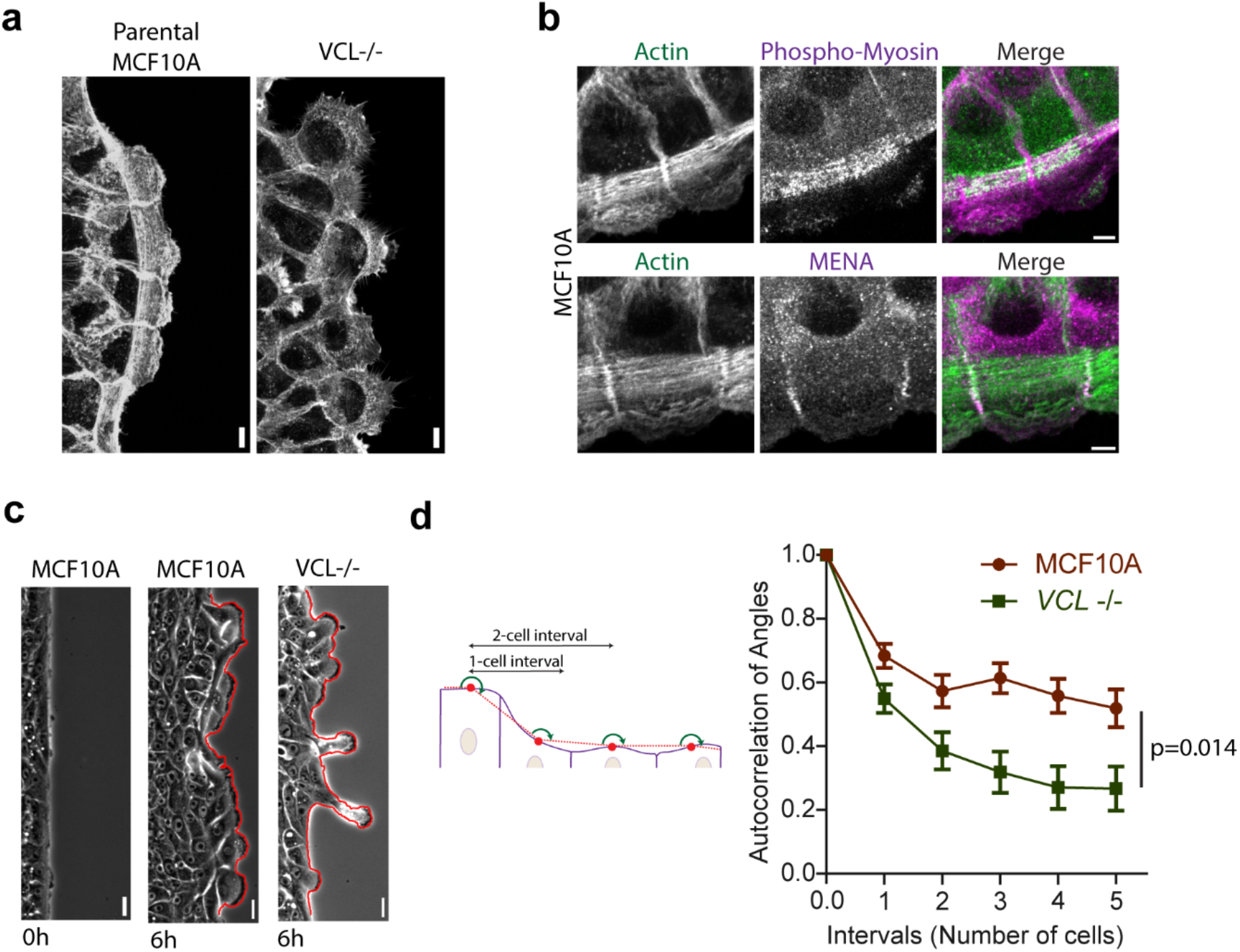
Vinculin coordinates lamellipodia during collective migration. **a** Staining of actin in MCF10A and VCL-/-cells 6h after migration of the leading edge. Scale bar = 5 μm. **b** Staining of actin, MENA and phospho-myosin light chain 2 (Thr18/Ser19) in MCF10A 6 h after migration of the leading edge. Scale bar = 5 μm. **c** Migration of a leading edge of MCF10A and VCL-/-cells imaged in phase contrast. Scale bar = 5 μm. **d** Persistence of the angles of the leading edge over multiple cell lengths. (Mean ± SD, One-way ANOVA of coefficient of exponential and plateau fits n=10, N=3. 1 representative experiment shown.)

**Figure 7:**
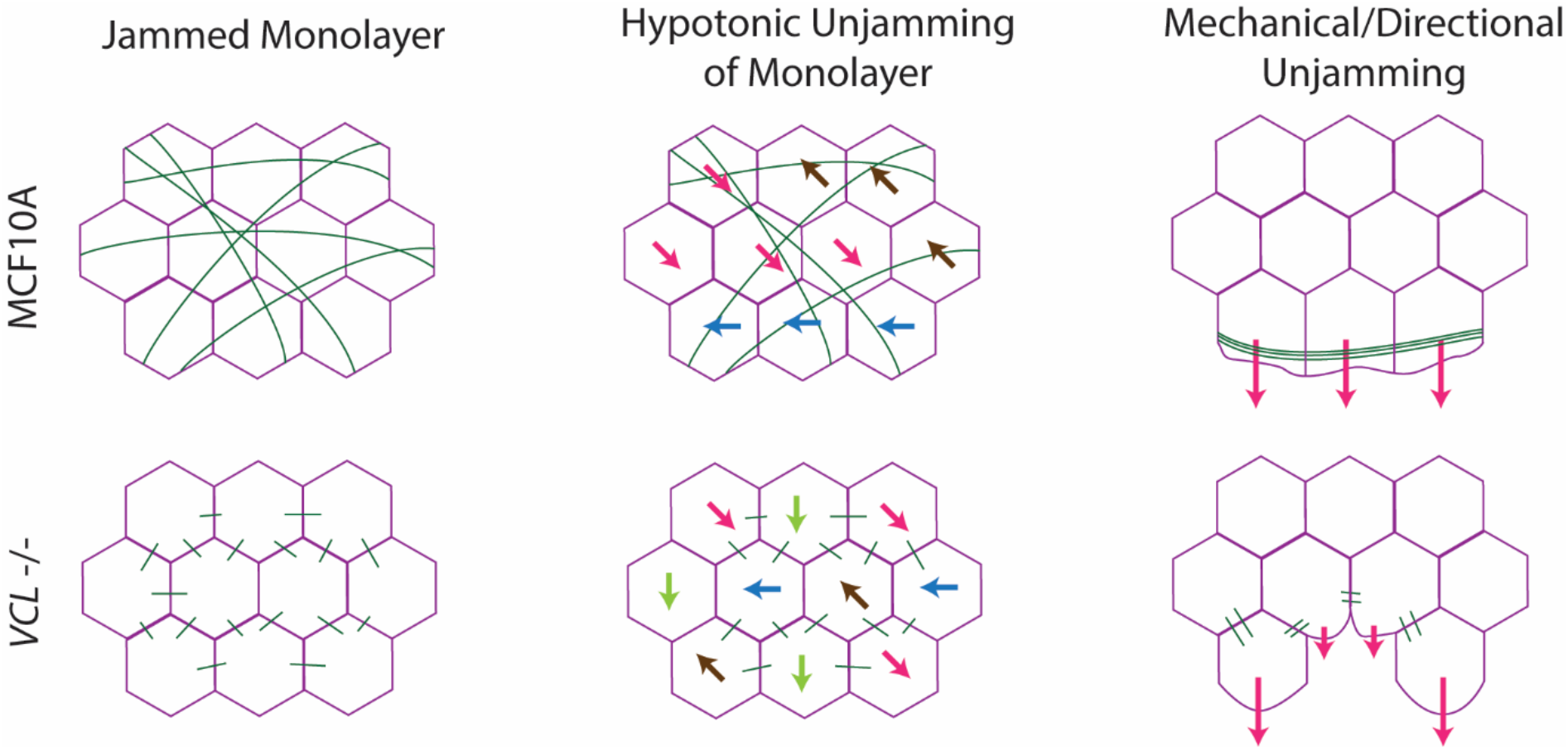
Long range TAFs lead to force anisotropy in MCF10A monolayers that lead to collective cell migration. Short range TAFs that connect adjacent uniformly in VCL-/-lead to random and uncoordinated cell migration.

MCF10A cells connected by TAFs seemed to be protruding lamellipodia collectively in the same direction (Fig 6a) while VCL-/-cells seemed to behave more independently creating lamellipodia in different directions. Thus, MCF10A cells create a more uniform edge of migration compared to VCL-/-cells (Fig 6c). To quantify this, we plotted a line along the leading edge, measured the angles made by the migrating front and analysed how well correlated these angles are over distance. This autocorrelation gives a measure of how collective the cells behave, with a maximum value of 1 indicating a perfectly uniform edge. We found that the autocorrelation curve decays over a shorter distance for VCL-/-cells compared to MCF10A cells, showing that the migration of MCF10A cells is correlated over a larger distance than that of VCL-/-cells (Fig 6d).

## Discussion

Here, we have shown that vinculin at a FAJs is essential for the coordination of TAFs across multiple cells. Cells connected by long-range TAFs migrate more collectively during unjamming by producing lamellipodia in a more coordinated manner. Our novel quantification of collectivity of migration in two different contexts of unjamming is unbiased by speed of migration and thus unbiased by the difference in ability of the cell lines to migrate individually. Hence, we believe that this shows the effect of vinculin at cell-cell adhesions on collective migration in isolation from the role of vinculin at cell-substrate adhesions.

TAFs are a pair of contractile actin bundles in adjacent cells connected by a focal adherens junction. Although similar structures have been previously reported in endothelial cells, we are the first to characterise their composition and formation in polarised epithelial MCF10A monolayers. We have shown that these fibres are formed at the apical pole of cells in mature monolayers, indicating they may be present in mature tissues. Perhaps most importantly, we show that they are formed in the physiologically more relevant mammary organoid system.

In a previous study, we had established that vinculin inhibits branched actin formation through its direct interaction with Arp2/3 in epithelial cells (James et al. 2025). Surprisingly, both upregulating and downregulating Arp2/3 globally in a cell leads to the loss of TAFs. Consistently, perturbing the regulation of Arp2/3 by vinculin, by abolishing vinculin-Arp2/3 binding using the VCL P878A mutation, also leads to the loss of TAFs. Exactly how Arp2/3 activity leads to the formation of TAFs remains unclear. One possibility is that Arp2/3 at early AJs could provide branched actin filaments that are bundled into TAFs by actin bundling proteins such as the vinculin tail.

Interestingly, our results also show that even though migration is driven by lamellipodia at the basal surface of the cells, AJs at the apical pole control the direction of migration. Our finding that anisotropic arrangement of TAFs, which can produce contractile forces, coincides with an increase in collectivity is in line with previous predictions that force anisotropy in a monolayer leads to collective migration. Additionally, during space-induced unjamming, we found that lamellipodia are produced and migration occurs perpendicular to TAFs. Pulling forces along the direction of a TAF from either side of the cell could inhibit migration perpendicular to the TAF. This suggests that the presence of a TAF constrains migrating cells into staying collective. This finding is supported by previous reports from wound healing experiments (Cochet-Escartin et al. 2014) in which cutting the actin fibre at the edge of the wound resulted in the emergence of a new leader cell, presumably from the loss of this mechanical constraint on migration.

It remains unclear why knocking out vinculin leads to TAFs becoming disorganised in the cell interior. It is possible that the organisation of the TAF network could be a result of large-scale self-organisation of contractile acto-myosin, where vinculin strengthening FAJs allows cells to better transmit forces across multiple cells in the monolayer. Another possibility is that vinculin, acting as a mechanosensitive hub capable of binding to several proteins, initiates a signalling cascade that organises actin fibres in the cell interior.

## Materials and Methods

### Cell culture, cell lines and reagents

MCF10A cells were cultured in DMEM:F12 GlutaMax (31331028, Gibco) supplemented with horse serum (5%, H1270, Sigma), EGF (20ng/ml,AF-100-15, Peprotech), hydrocortisone (0.5ug/ml, H0888-1G,Sigma), insulin(0.1%, I9278, Sigma), cholera toxin(0.1ug/ml, C8052, Sigma), Pen-Strep(1%, 15140122, Gibco). Cells were trypsinised (12605010, Gibco) and subcultured every 3 days. VCL-/- and VCL P878A-KI cell lines were generated by CrispR-Cas9-based genome editing (James et al., 2025), and RacQ61L was stably expressed (Molinie 2019). For CK666 treatment, cells were allowed to adhere for 6 hours in media without CK666 before adding the drug.

### Immunofluorescent staining and antibodies

Cells were fixed with 2% PFA and permeabilized for 5 min in 100% ethanol at -20°C, followed by blocking in 10% FBS in PBS. Primary antibodies against vinculin (#V9131, Sigma), E-cadherin (#MABT26, Merck), MENA (mAb A351F7D9), phospho-myosin light chain 2 (Thr18/Ser19) (#3674, Cell Signalling) or laminin (#MAB19562,Sigma) were used followed by Acti-Stain 555 (Cytoskeleton) for actin, along with secondary antibodies, anti-mouse-647 (#A21236, Life Technologies) and anti-rabbit-405(#A34556, Life Technologies) or anti-rat-647(#A11007, Life Technologies). Coverslips were mounted in Dako mounting medium and imaged the next day.

For growing acini, cells were seeded on top of polymerized Matrigel (CB-40230C, Corning) in Millicell EZ SLIDE 8-well glass chamber slide (PEZGS0816, Millipore) in a medium containing 4ng/mL EGF (4ng/mL) and 1% serum and supplemented with 2% of Matrigel. The medium was changed every 3 days and acini were allowed to grow for 3 weeks. Acini were fixed in 2% PFA in PBS and permeabilized with 0.5% Triton X-100. Acini were then rinsed in PBS with 100mM glycine and blocked successively with IF Buffer (PBS with 10% FBS, 0.1% BSA, 0.2% Triton X-100, 0.05% Tween-20) and IF Buffer + 20 µg/ml goat anti-Rabbit Fc fragment (111-005-046, Jackson ImmunoResearch). Acini were incubated with the primary antibodies followed by IF buffer washes and then incubated with secondary antibodies and phalloidin. Slides were mounted with Abberior Mount Liquid Antifade (Abberior) and sealed with valap. Images were acquired with a 40x water immersion objective on a Leica SP8 laser-scanning confocal microscope.

### Imaging and Analysis

For analysing the migration of the leading edge, 1.5×10^6^ cells cm^-2^ were plated on Ibidi dishes (#80466) with inserts. Inserts were removed 1 day after plating and phase contrast images were acquired on an Olympus IX83 microscope using a 20x objective (NA 0.5) equipped with an Orca-Flash4.0 V3 camera (Hamamatsu). Coordinates of the leading edge of migration were extracted using the FIJI plugin JFilament. Analysis of the directional persistence of the leading edge over distance was done using algorithms previously described for analysing persistence of angle of migration of cells over time (Gorelik and Gautreau 2014). For hypotonic unjamming, 1×10^6^ cells cm^-2^ were plated on ibidi Ph+ microslides (#80446) and treated with hypotonic medium (9 parts cell culture medium and 1 part distilled water) after 3 days. Unjamming cells were imaged at 10-min intervals on an Olympus IX83 microscope using a 20x objective. PIV analysis was carried out as described in (Garcia et al. 2015).

For imaging the transcellular actin network, 1.5×10^6^ cells cm^-2^ were plated on fibronectin (10 µg/mL, F1141, Sigma) coated coverslips and stained 3 days later. Z-stacks were taken at 0.2 μm intervals using a Leica SP8 laser-scanning confocal microscope, and stacks were deconvolved using an adaptive blind algorithm based on theoretical PSFs in Autoquant X2 (Mediacybernetics). Number and percentages of cells crossed by TAFs were counted manually in FIJI.

## Author contributions

J.J. performed all the experiments and analyses, and wrote the first version of the manuscript. A.M.G and S.R. have jointly supervised the work and S.R. wrote the finale version of the manuscript.

